# Nuanced patterns of biological collecting trends revealed by informatic investigation of zoological natural history collection records

**DOI:** 10.1101/2023.09.29.560208

**Authors:** Daren C. Card

## Abstract

Natural history museums comprise a unique and important component of biological infrastructure worldwide, and underlie diverse research, education, and outreach in the natural sciences. Each museum is built around one or more biological collections that serve as a repository of important materials for study, which have contributed to scientific research in increasingly important areas ranging from understanding global climate change to developing new biotechnological applications. However, despite centuries of existence, global collections sizes have only been recently estimated, and aside from analyses of certain institutions or well-studied clades, little is known about how patterns of collecting activity across institutions, geographic regions, or taxonomic clades, or how these patterns vary across time. To address this important gap in our understanding of critical life sciences infrastructure, I gathered and analyzed zoological records associated with preserved specimens collected between 1900 and 2015 and housed in worldwide natural history collections using data from the Global Biodiversity Information Facility. My analysis indicated that global, museum-associated collecting activity focused on animals has varied greatly over time, peaking around 2000 and declining significantly since. By stratifying data by institution, nation, and taxonomic phylum and class, I also illuminated how individual data series contribute to these global patterns. Institutions and nations had either concentrated or dispersed periods of relatively high collecting activity that occurred over different time periods, although most growth occurred in the second half of the 20th century for both many individual institutions and globally. Certain taxonomic clades comprise the largest proportions of collections records over this time period, namely arthropods (especially insects) and chordates (especially vertebrates), underscoring the taxonomically biased collecting histories of many institutions. Altogether, my analyses provide a critical, early view of historical zoological collecting activity and, to help other museum stakeholders to explore the data and results in more depth, I also describe and release NHMinformatics, an interactive dashboard built using R Shiny. These resources, when combined with recommendations I have made for sustainable collections growth, will be helpful for establishing policy goals, provisioning museum infrastructure, and training curatorial personnel.

## Introduction

Natural history museums are important institutions in the life and earth sciences and are vital cultural centers with a worldwide distribution. Their primary mandate – largely established when advances in seafaring technology catalyzed worldwide exploration and the need to build repositories to store materials for future study – is to house valuable specimens and artifacts collected from across the globe. Therefore, natural history museums are leading institutions of biodiversity research, a role that has taken on increased purpose with the ongoing biodiversity crisis. However, these institutions also form a foundation for diverse research areas in biology, geology, and other disciplines, but have also adopted the mission of championing education and public outreach in the natural sciences. Research and education in natural history museums has been broadly influential in addressing diverse societal issues, including combating global change, human disease, and food insecurity, and bolstering environmental conservation, sustainable harvest of natural resources, and the emerging bioeconomy . More directly, research leveraging natural history museums has illuminated the link between El Niño-driven precipitation, increased density of rodents in the US Southwest, and heightened risk for human contraction of Hantavirus Pulmonary Syndrome in this region (Yates et al. 2002). Natural history collections have also helped elucidate the impact of a century of climate change on altitudinal distributions and ecologically-relevant genetic diversity of small mammal populations in Yosemite National Park, California, USA (Moritz et al. 2008; Bi et al. 2019). Moreover, more distant fields, such as engineering, have also been influenced by natural history collections, as solutions for noise reduction in Japanese high-speed trains has been inspired by owl and kingfisher morphology (McKeag 2012).

Considering the importance of natural history museums for modern research, education, and outreach, one would expect that museum holdings would be enumerated and well understood. However, unfortunately, a complete accounting of the collections of natural history museums does not exist, leaving scientists with largely anecdotal information about the sizes and compositions of the world’s natural history museums. Only recently have estimates of worldwide collection holdings been amassed, after years of dedicated collection digitization that have resulted in only about 30% of U.S. natural history samples being digitized and made available in electronic databases (National Academies of Sciences, Engineering, and Medicine 2020). Since 2004, the Global Biodiversity Information Facility (GBIF) has aggregated billions of organism occurrence records, including, as of 2023, over 200 million occurrences linked to preserved specimens stored in natural history museums, herbaria, and other biorepositories (GBIF.org 2023). Digitization of natural history museum holdings has generally advanced most greatly in the United States, which is due in part to governmental investments through the National Science Foundation Advancing Digitization of Biodiversity Collections (NSF ADBC; https://www.nsf.gov/funding/pgm_summ.jsp?pims_id=503559) program, where the central coordinating organization iDigBio has coordinated the aggregation of a large proportion of the data included in GBIF (Nelson & Ellis 2018). Recently, a targeted survey of 73 of the world’s largest natural history museums and herbaria located in 28 countries determined that there are more than 1.1 billion specimens or artifacts in natural history collections, which were composed of 19 collection types sourced from 16 global geographic areas (Johnson et al. 2023). While significant work remains in digitizing and integrating natural history collections data worldwide, the survey by Johnson et al. (2023) is the first to use full data from the largest museums to estimate the minimum size of worldwide collections holdings, making it critically important for deciding on future research funding levels, infrastructure investments, and workforce development needs.

Estimates of the sizes of natural history museum collections are valuable for creating policy that helps ensure that the infrastructure and personnel needs of natural history museums are being met. However, collection size data comprises contributions to natural history museums over their entire history, providing relatively coarse understanding of overall collections activity. Estimates of collection sizes are valuable for understanding the scale and cost of curating existing collections into the future but provides essentially no information about temporal patterns of collecting activities across various institutions and taxonomic groups, or how such activity may continue. Information on temporal collecting activity over the combined and individual histories of natural history museums provides a much richer picture of an institution’s status that is theoretically more useful for making policy decisions. For example, funding needs may differ between natural history museums that maintain high collecting activity and growth versus historically more active institutions that are now growing much more slowly, or even between collections with different taxonomic or geographic foci with distinct collection and curation considerations. Moreover, understanding global patterns of collections growth across natural history collections is vital for assessing the health of natural history museum collections, which would ideally have sustainable collecting activity over time, and the utility of these resources for addressing a range of biological questions.

While data-informed estimates of the sizes of natural history museum collections are beginning to emerge, there is little publicly available information on the sizes of natural history collections that can be used to understand temporal patterns of collecting activity and museum growth. Using data from GBIF for vertebrates, Rohwer et al. (2022) found declines in the deposition of vertebrate specimens to natural history collections since peaks in collection activity that occurred in the 1970s and 1980s, depending on the taxonomic class. Collecting trends in Western Hemisphere mammal collections have been periodically assessed by the American Society of Mammalogists, most recently in 2018, with noticeable declines in collecting activity observed in recent years (Dunnum et al. 2018; Malaney & Cook 2018; Cook & Light 2019). In contrast, a recent evaluation of collections holdings and digitization efforts of North American arthropod collections found that the number of specimens in collections has increased by approximately 1% annually over 30 years, although the authors note that this rate of increase is insufficient for addressing many biodiversity research needs (Cobb et al. 2019). Moreover, data from five major U.S. natural history museums indicated recent growth of cryogenetic collections, a more recent form of collection that includes important materials for molecular investigations comprising the emerging field of museum genomics (Card et al. 2021). However, for most natural history museums, individual collections, time periods, and taxonomic groups, information on collecting activity and collection growth is lacking, leaving natural history collections poorly equipped to assess curatorial costs of existing collections, predict future collecting targets and trends, and advocate for sustainable levels of financial and infrastructure support. Anecdotal evidence also suggests that collecting activity at natural history museums has declined in recent years, although no published studies have confirmed whether this is true.

To begin addressing this knowledge gap, I filtered and standardized global, publicly available records for animals derived from preserved specimens in natural history collections. My focus on animals alone was due to my taxonomic expertise with this clade and to computational and analysis constraints stemming from these large datasets, but my approach could also be applied to data from plants or fungi. Based on these data, I evaluate historical patterns of collecting activity across well-represented taxa in the largest natural history museums with catalog data available through GBIF. I also describe a new informatics dashboard built upon these filtered, standardized museum specimen records that will facilitate additional investigations of these data by institutions and individuals across the globe. After characterizing global collection trends and the temporal accumulation of zoological specimens across different nations, institutions, and taxonomic units, I conclude with several recommendations for ensuring sustainable growth and general vitality of natural history museum collections.

## Materials & Methods

Publicly available records for animals deposited into natural history collections were retrieved from GBIF using the options “Basis of record” = “Preserved specimen” (excludes fossil specimens) and “Scientific name” = “Animalia” (N = 99,513,214 records; retrieved 2023-01-31; https://doi.org/10.15468/dl.uxhkny). These records represent the subset of natural history museum holdings that have been digitized and required filtering and standardization for comparison and visualization. I filtered the records as follows: (1) records where the Kingdom was not ‘Animalia’ were removed; (2) records not identified to the level of genus or below were removed; and (3) records with missing or undefined data for year collected, institution code, or collection catalog number were removed (N = 61,248,185 records retained). I summarized the number of records for each taxonomic phylum and class, collection code, and year, associated each individual collection code with a parent institution or publisher (i.e., some institutions publish subcollections data to GBIF separately), and focused on records with a defined taxonomic phylum and class that were collected between 1900 and 2015. To avoid potential biases in taxonomic clades and institutions with small numbers of records, I focused on well-represented phyla and institutions by selecting phyla with at least 5,000 records worldwide and institutions with a minimum of 100,000 records amassed between 1900 and 2015. I also ensured that phyla were consistently collected between 1900 and 2015 (at least 100 records per year for at least 50 years total) and institutions had relatively consistent collecting activity between 1900 and 2015 (at least 100 records per year for at least 50 years total). 66.6% of records (N = 40,816,928) were retained after applying these filters. Given that the number of records for collections and taxonomic groups can differ significantly, making comparison and visualization difficult, I standardized the number of records for each institution and year by the total number of records amassed per institution between 1900 and 2015, producing an annual measure of collecting activity that is a proportion of the institutional collection growth over the 1900 to 2015 timespan. Using the resulting dataset, I assessed overall global collections growth and temporal, institutional, and taxonomic patterns of collecting activity between 1900 and 2015. I also used Shiny v. 1.7.4 (Chang et al. 2023) in R v. 4.2.3 (R Core Team 2023) to build a supplemental interactive dashboard that will allow others to explore natural history collections growth.

## Results

The 40,816,928 zoological records surviving quality filtering comprised 72 taxonomic classes and 10 phyla and represent the holdings of 102 institutions from 21 countries and all inhabited continents (Australia was included with Oceania). Although the total and relative size of natural history museums was specifically addressed in Johnson et al. (2023), my data provided similar, although less complete, information on the relative sizes of the largest institutions, which were concentrated in Europe and, especially, North America. In the following sections, I briefly describe major patterns of historical collecting activity.

### Global zoological collections growth has peaked

The most glaring and unfortunate result of investigating historical patterns of zoological collection growth is that global zoological collections growth has peaked and currently appears to be in steady decline (Figure 1). Collecting activity was relatively modest in the early decades of the 20th century before noticeable growth beginning around 1920 that continued through the late 1930s. World War II had a marked, negative impact on collecting activity in the 1940s and it was not until the around 1950 that collections growth matched pre-war levels. Beginning in the late 1950s and continuing until about 1965, there was significant growth of natural history collections. Growth then plateaued or modestly declined through the late 1980s before another era of substantial growth that extended through the early 2000s, which represents the period when collecting activity and growth of natural history collections globally was greatest. Since this peak, activity has declined precipitously – at a rate similar to declines observed during World War II – through 2015, the final year evaluated (Figure 1). Activity likely remains diminished to date, although I avoided assessing patterns after 2015 due to potentially incomplete data from recent years. Annual collections growth in 2015, and presumedly today, matches growth in around 1960, which is the approximate start of an era of unprecedented growth during the second half of the 20th century. Therefore, investments and concerted effort are needed to prevent collections growth from falling to levels not seen since before the modern era of science research that began in the wake of World War II.

**Figure 1.**
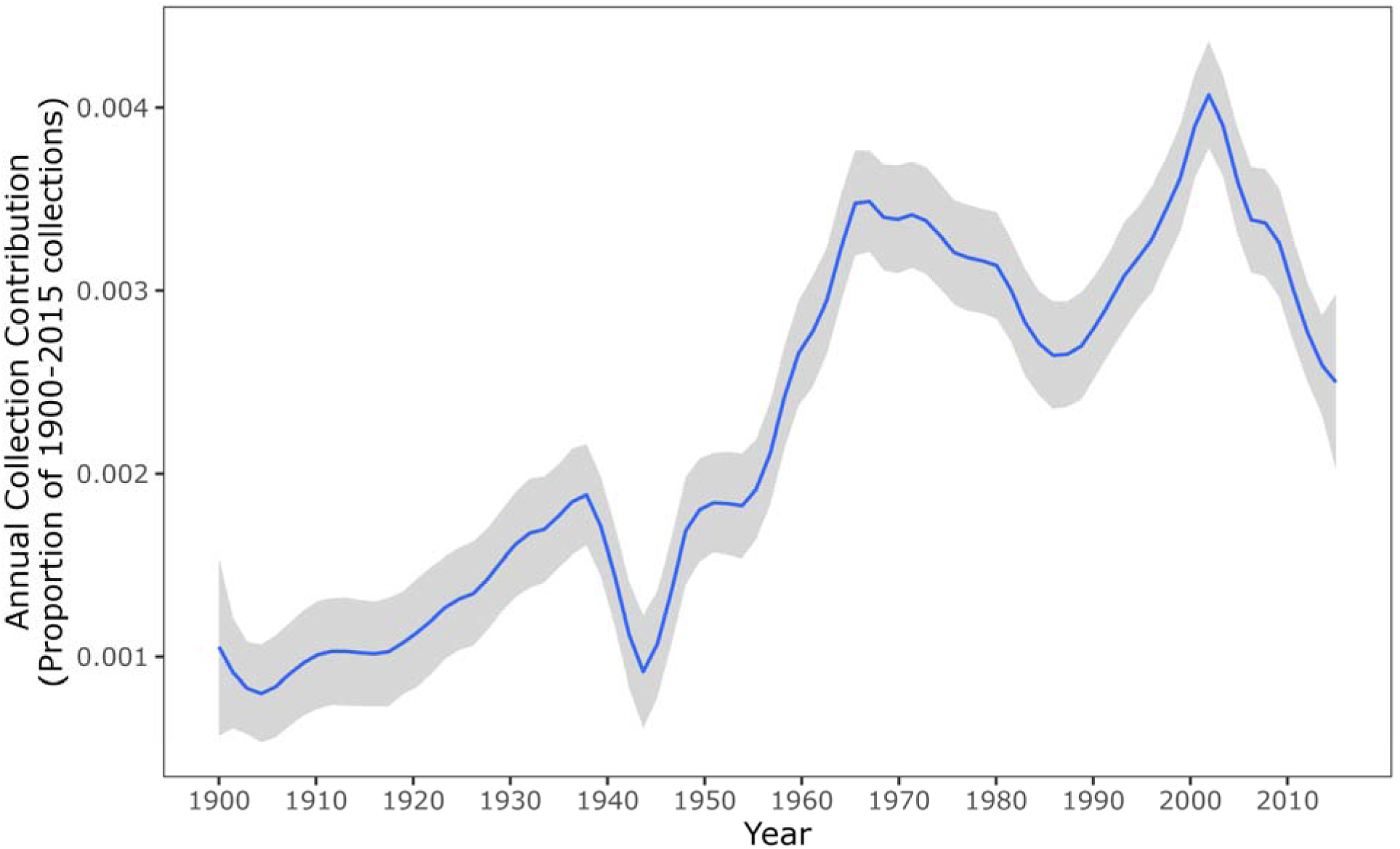
Assessment of historical growth of natural history collections indicates a peak and subsequent decline in collecting activity. Temporal patterns of the number of specimens deposited in worldwide natural history collections between 1900 and 2015, as based on data from the Global Biodiversity Information Facility (GBIF). Annual activity is reported as a percentage of total collections made across all clades and institutions over the entire time period after rigorous data filtering.

### Domain-specific patterns of collections growth provides nuanced understanding of collecting activity

In much the same way that temporal patterns of collecting activity provides richer information than total collection sizes alone, assessing patterns of collections growth for individual taxonomic units, natural history museums, and nations leads to even deeper understanding of the health and vitality of global biodiversity collection activity and the biorepositories where specimens are stored. My analysis of taxonomic, national, and institutional patterns of zoological collecting activity between 1900 and 2015 indicates biased accumulation of taxa and varying national and institutional growth patterns. At the level of phylum, standardized collecting activity indicates that most taxa have been most heavily collected since 1950 (Figure 2A). Based on raw number of records, Arthropoda and Chordata are by far the most collected phyla (at least 100,000 records for each 5-year period since 1900), although Mollusca, Cnidaria, Echinodermata, and Annelida have also been collected in relatively high numbers (at least 10,000 records for each 5-year period since ∼1960; Figure 3A). Porifera, specifically, shows a different pattern where collecting activity is concentrated in the period since 1980. Collecting patterns in Arthropoda and Chordata drive much of the overall temporal trends across taxa (∼55% and ∼34% of all records, respectively), and within these phyla, certain taxonomic classes, namely Insecta and various vertebrates, represent most of the collecting activity (Figure 2A and Figure 3A). Collecting activity focused on Chordata peaked in the 1960s and was relatively healthy into the 1990s, but has since waned significantly, especially in the case of amphibians and reptiles, mirroring vertebrate-specific results presented in Rohwer et al. (2022). Vertebrates are the most collected chordates and the collecting trends of each vertebrate taxonomic class generally mirror overall patterns in Chordata with high collecting activity in the 1960s through the 1990s, except in the case of Aves, which were collected in relatively high numbers only in the 1960s, but also exhibited a burst in collecting activity far earlier than other vertebrates (1905-1940; Figure 2A). Aves is also the only taxonomic class outside of Insecta where at least 100,000 specimens were collected during each 5-year period between 1900 and 2015 (Figure 3A), although consistently large collections of mammals have accumulated since 1930 and gastropod molluscs have been well represented in collections since 1960. In contrast, collecting activity in arthropods has continued to grow over time, but has leveled off in recent years. Consistent collections of insects dominate these trends, although Malacostraca was relatively consistently collected between 1900 and 2015 and Arachnida and Copepoda are characterized by relatively high collecting activity after 1990 (Figure 3A). Indeed, if not for significant growth in collecting activity in Arthropoda beginning around 1990, which is well justified by the proportions of known and unknown biodiversity in this clade, overall collections growth likely would have peaked much earlier in the 1960s and the subsequent decline in activity would have been far more precipitous (Figures 1 and 2A).

**Figure 2.**
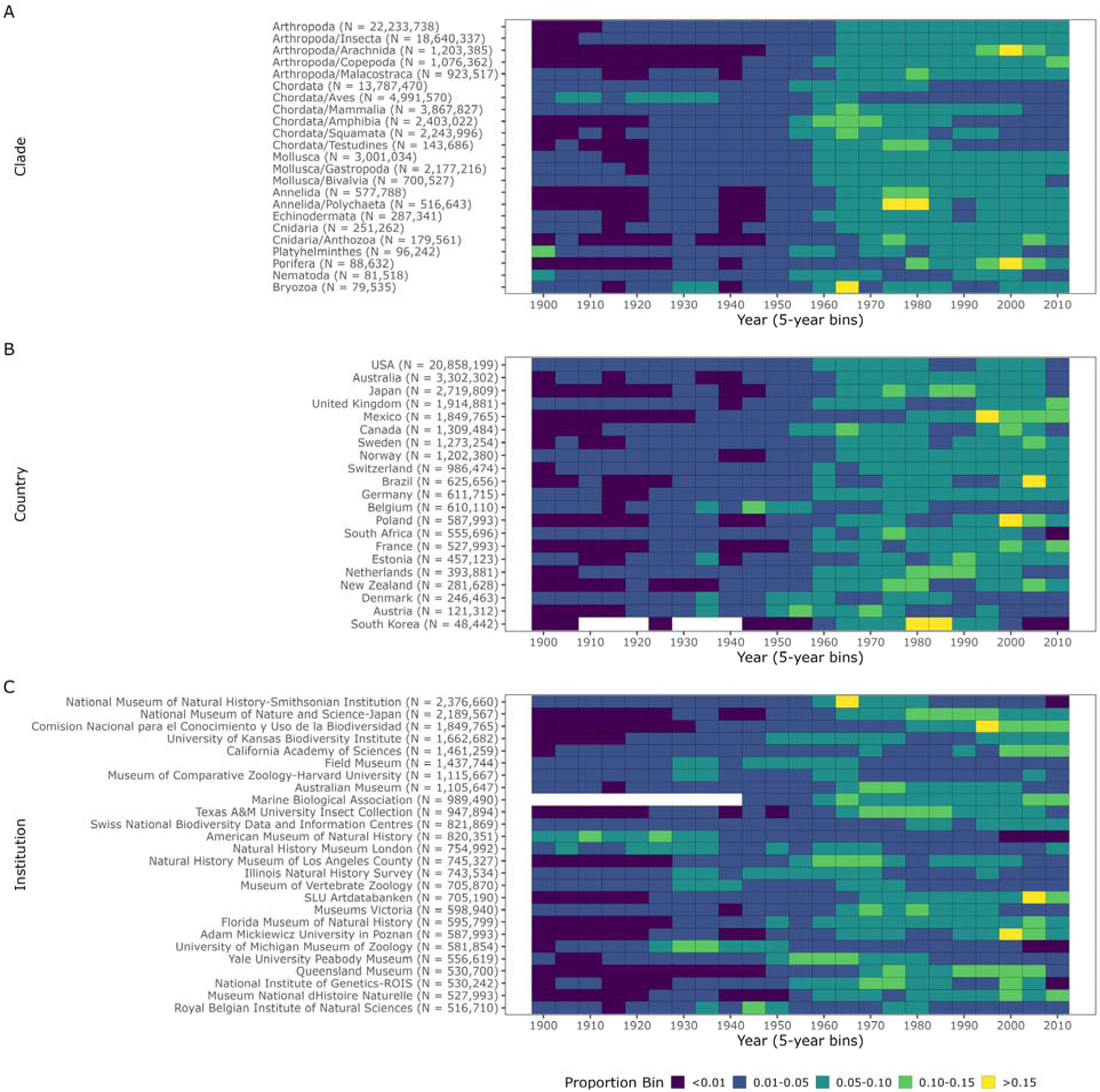
Idiosyncratic patterns of historical natural history collections growth across taxonomic clades, nations, and natural history museums/institutions. Sliding window measures of standardized collecting activity over 5-year intervals for (**A**) consistently-collected taxonomic phyla (see above) and classes (at least 100,000 global records) as a percentage of total global collections for each clade between 1900 and 2015, (**B**) 21 nations with at least one museum/institution as a percentage of total collections for each nation between 1900 and 2015, and (**C**) 26 of the largest natural history museums/institutions with total collection sizes greater than 500,000 records each as a percentage of total collections for each museum/institution between 1900 and 2015.

**Figure 3.**
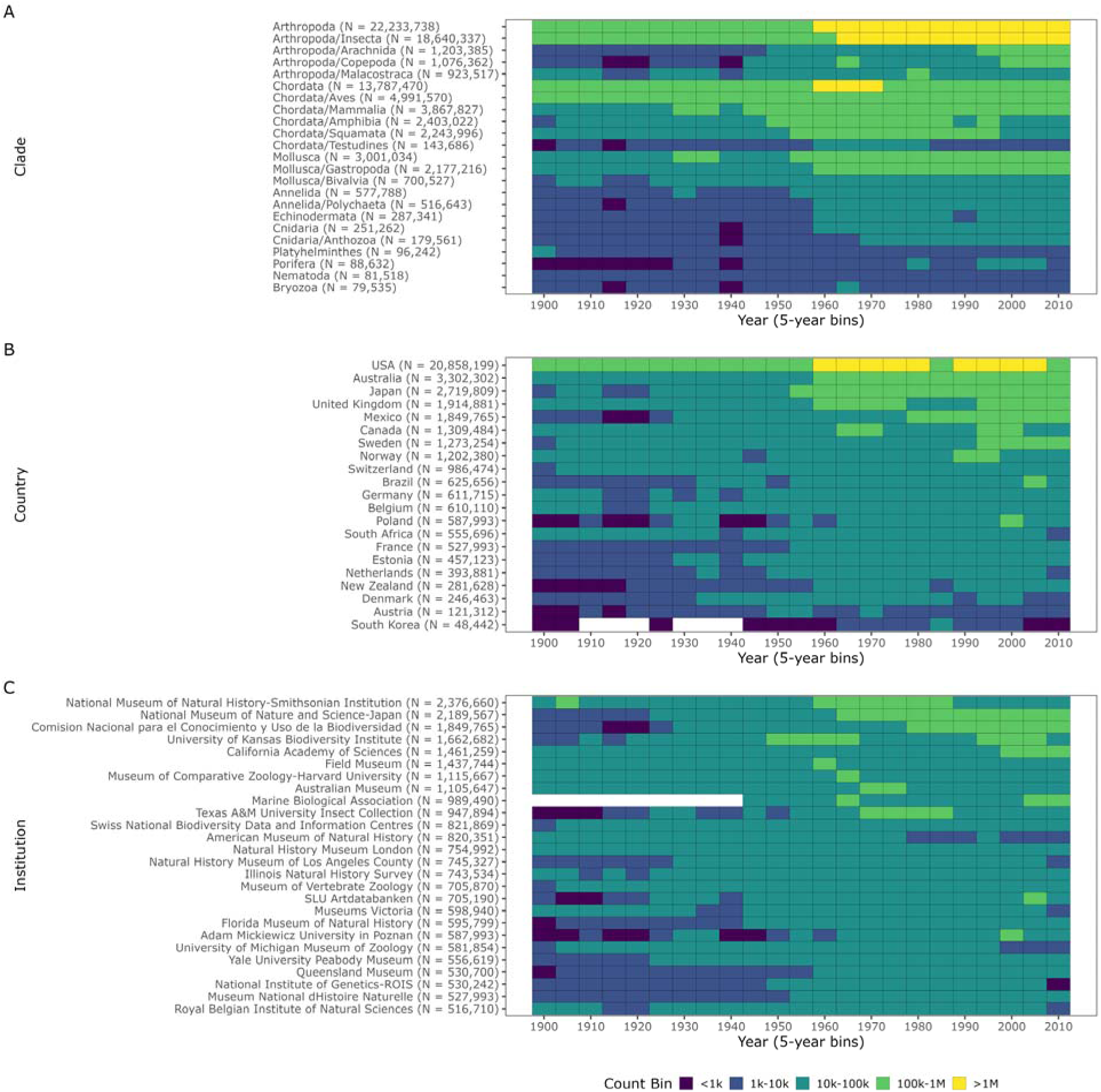
Variable growth of natural history collections records for different taxonomic clades, nations, and natural history museums/institutions. Sliding window measures of collecting activity over 5-year intervals based on absolute counts of records of (**A**) consistently-collected taxonomic phyla (see above) and classes (at least 100,000 global records), (**B**) 21 nations with at least one museum/institution, and (**C**) 26 of the largest natural history museums/institutions with total collection sizes greater than 500,000 records each.

Given that national zoological collecting trends, in most cases, integrate over multiple collections, they are relatively homogenous and most records have accumulated since about 1960, although several European nations (Austria, Belgium, Denmark, and Estonia) had bursts of relatively high collecting activity between 1920 and 1960. In contrast, collecting activity in Mexico has been concentrated in the period since 1980 (Figure 2B). Collections growth patterns in the United States dominate overall trends due to the concentration of large natural history museums (∼51.5% of records are from U.S. museums), which reflects the concentration of major natural history museums in the United States and early digitization efforts through NSF ADBC. At least 100,000 records have been amassed every 5-year period since 1900 and since 1960 and the U.S. has accumulated more than 1 million records over most 5-year periods. Only Australia, Japan, the United Kingdom, Mexico, Canada, and Sweden have had multi-decade sustained growth of collections exceeding 100,000 records per 5-year period, and Norway, Brazil, and Poland have also had periods of relatively high, though less sustained, collections growth (Figure 3B). On the other hand, individual institutions show more marked differences in the accumulation of natural history records. Collecting activity was relatively high before 1950 for certain collections, such as the American Museum of Natural History, Natural History Museum London, University of Michigan Museum of Vertebrates, Field Museum, Museum of Comparative Zoology at Harvard University, Illinois Natural History Survey, Museum of Vertebrate Zoology, and Royal Belgian Institute of Natural Sciences (Figure 2C). Most other institutions had periods of relatively high collecting activity that began in the 1950s and, especially, the 1960s, which corresponded with the significant overall growth of natural history collections in this period. For many of these institutions, these periods of relatively high collecting activity have persisted beyond 1970 and, in some cases, to the final period evaluated in this study, such as the National Museum of Nature and Science – Japan, University of Kansas Biodiversity Institute, the Marine Biological Association, the Florida Museum of Natural History, Adam Mickiewicz University in Poznan, and the Muséum National d’Histoire Naturelle (Figure 2C). Other institutions – namely the Comisión National para el Conocimientio y Uso de la Biodiversidad (Mexico), California Academy of Sciences, Swiss National Biodiversity Data and Information Centres, SLU Artdatabanken, and Queensland Museum – have had most of their collection growth occur since 1970 (Figure 2C). However, based on raw record counts, only a small number of the largest institutions had relatively consistent, high collecting activity (at least 100,000 records) for at least two 5-year periods between 1900 and 2015: National Museum of Natural History – Smithsonian Institute, National Museum of Nature and Science Japan, Comisión National para el Conocimientio y Uso de la Biodiversidad (Mexico), University of Kansas Biodiversity Institute, California Academy of Sciences, Australian Museum, Marine Biological Association, and Texas A&M University Insect Collection (Figure 3C). National and institutional collection trends underscore how variable zoological collecting activity can be and provide richer understanding of the health of global biorepositories individually and collectively.

### An interactive R Shiny application for exploring patterns of collections growth

The FAIR Data Principles (Wilkinson et al. 2016) of ensuring that data are **F**indable, **A**ccessible, **I**nteroperable, and **R**eusable are increasingly important in the current era of “big data”. Although natural history museums remain incompletely digitized (National Academies of Sciences, Engineering, and Medicine 2020) and have only recently begun to integrate data summarizing their vast collective holdings (e.g., (Johnson et al. 2023)), in many ways, these institutions were one of the pioneers of the FAIR Data Principles, at least in the non-digital realm. To continue this tradition and following the recent example of Johnson et al. (2023), I used R Shiny to build a simple, interactive application that will allow anyone to explore the filtered, standardized data that have been summarized in this article. This application has been provided as an R Markdown document and contains a summary of the filtered, standardized data and embedded Shiny elements that facilitate interactive exploration of these data according to user interests. The document also includes a detailed supplement with the full details – including embedded code, plots, and tables – of retrieving, filtering, and standardizing the raw GBIF data, an overview of the full exploratory analyses and visualization that formed the basis for this investigation, and all code and documentation necessary to reproduce the analyses and figures presented in this article. The resources produced as part this article have been made permanently available through a Zenodo-archived repository (https://doi.org/10.5281/zenodo.8393331), which contains a partially filtered intermediate dataset used for data summarization and exploration in R, a final dataset summarizing the records of different taxonomic groups and institutions, and an R Markdown document with embedded Shiny elements that enables user exploration of the data. In the spirit of FAIR, interested users will always be able to download the permanently archived repository and use R to run the R Markdown document to interactively explore the data on their local computer. Additionally, instructions for accessing an internet-hosted version of the R Markdown document are available at https://github.com/darencard/NHMinformatics to allow interested users to explore the collections data more easily. Any future updates to these resources will be made available through the Zenodo-linked GitHub repository (https://github.com/darencard/NHMinformatics).

## Discussion

Using data from over 40 million cataloged zoological records from global natural history museums, I provided the first published summary of temporal collections growth based on records from 10 well-collected phyla and the 102 largest institutions who have contributed data to GBIF. The growth of these natural history collections has varied significantly over time, reflecting distinct eras of global natural history collecting activity. Beginning in 1900, the first year evaluated in this study, natural history museums saw approximately 70 years of consistent growth only interrupted by a sharp downturn and subsequent recovery during and after World War II. This pattern is a marked departure from the trends observed since around 1970, and following the peak of collections growth in 2000-2001, collecting activity has fallen precipitously. Variable patterns of collecting activity across nations, institutions, and taxonomic clades underly these complex temporal trends and highlight the unique contributions individual institutions have made to global natural history museum holdings. Inference of these patterns relied upon rigorous filtering and standardization that resulted in over 40 million records, encompassing institutions that are known to be among the largest in the world, which provides confidence that the true temporal patterns in global collecting activity have been captured.

While great care was taken in filtering and standardizing collections data, these data had inherent biases from the start and decisions during the analysis process may have introduced further biases that are difficult to identify and overcome. Varying proportions of collections data have been excluded from the analysis due to filtering decisions (approximately 60% across the full dataset) and therefore, collections data from some well-known major collections were not included or were only included in part in the analysis. For example, most records (95%) from Museum of Vertebrate Zoology were included in the final analysis while only 58% and 25% of National Museum of Natural History-Smithsonian Institution and Natural History Museum London were included, respectfully, highlighting differences in data quality and standardization that impacted this investigation. As has already been noted, different countries and their individual institutions are in different stages of digitizing natural history collections, so the completeness of records in these cases may have impacted the results of this investigation, especially in cases where major collections remain largely “dark” because of non-existent digitization efforts. Moreover, even for well-digitized institutions, the pace and order in which institutions catalog specimens and digitize collections data can also vary, and even within an institution, different collections can have significantly different practices. For instance, some collections may prioritize digitizing older specimens due to their increasing importance in temporal investigations in an era of global change while other collections may give priority to newer specimens due to higher material quality or ease in gathering more-recently collected metadata. Due to differences in collecting and curation practices certain taxa may be prone to more biases. For example, fishes, the most biodiverse chordate group, are conspicuously absent from Figures 2-3 despite being well collected, a pattern that could be due to a range of factors discussed here and also due to the practice of collecting and cataloging fishes in lots and not individually (Hilton et al. 2021). Even the composition of individual personnel at institutions can be impactful, as an especially active collector or an individual with international expertise in taxonomy can lead to temporary spikes in collecting activity. My decision to focus on records identified to the level of genus leaves my analysis prone to the influence of a taxonomic expert, especially in the case of understudied or mega-diverse taxa, and this may explain pre-1950 spikes in the collection of Platyhelminthes, Nematoda, and Bryozoa (Figure 2). However, in restricting my analysis to consistently collected taxa from institutions with consistent collecting activity between 1900 and 2015 and by focusing on global patterns of high-level taxa (phyla and classes), my analysis is more resilient in the face of potential biases. Indeed, the parallel patterns of post-World War II growth and recent decline across institutions, nations, and major clades suggests the result of recent declining collecting growth is real despite any underlying biases. Factors that could confound this analysis must be identified so that data can be collected via international surveys of biological collections to enable a properly controlled, model-based investigation of this important question. Fortunately, as part of recent efforts to integrate worldwide natural history collections data, Johnson et al. (2023) also describe efforts to survey the size and age distribution of the museum workforce, gathering information that will be invaluable for future investigations.

The world’s largest 73 natural history museums and herbaria house more than 1.1 billion specimens and artifacts (including plant specimens, fossils, and cultural artifacts not assessed here; (Johnson et al. 2023)), reflecting the incomplete availability of natural history collection catalog data in GBIF, which suggests that further data and research are needed to precisely understand global collections growth, as roughly 4% of these holdings were included in my analysis. Although interesting for a variety of reasons, I also ignored the source locations for individual records when aggregating the data due to difficulties validating coordinates, inferring coordinates based on locality descriptions, and associating records with a nation or region. But anecdotal information and published data (Johnson et al. 2023) indicate that the source nations of museum specimens are less concentrated and geographically biased than the institutions that house them, which stems largely from historical practices during the colonial era that persist today, presenting unfortunate obstacles for nations and populations wishing to derive cultural, intellectual, and economic benefits from their natural heritage (Sheets-Pyenson 1987; Das & Lowe 2018; Ashby & Machin 2021; Nicolaï 2022). Overall, given that GBIF represents science’s best effort towards natural history museum collection digitization and integration, this research provides a meaningful, early representation of these important data trends that will form an important foundation for future collection data integration and analysis. Indeed, by making an interactive R Shiny application available for others to interactively explore museum collections growth trends, I continue a long tradition in natural history museum science of making data or materials publicly available for reuse. Although it is now possible to summarize specimen collecting trends across individual countries, institutions, and taxonomic clades – which should assist individual institutions and government institutions or interest groups in defining strategic collections priorities – the global trend of declining contributions to natural history museum collections uncovered in this study is concerning and undermines the mission of these important institutions.

The reasons for an apparent decline in biological collecting activity are multifaceted and the contribution of each factor to this pattern is currently poorly understood. Changes in regulations have undoubtedly negatively impacted collecting activity. Internationally, the Convention on the International Trade in Endangered Species of Wild Fauna and Flora (CITES; https://cites.org/eng), which was adopted widely in 1975, restricts the international trade of specimens of wild animals and plants, creating bureaucratic obstacles to scientific collecting that may contribute to declining collecting activity. The more-recent Convention on Biological Diversity, which went into effect in 1993 (Secretariat of the Convention on Biological Diversity 2000), and Nagoya Protocol on Access to Genetic Resources and the Fair and Equitable Sharing of Benefits Arising from their Utilization to the Convention on Biological Diversity (Secretariat of the Convention on Biological Diversity 2011), which entered into force in 2014, are additional international agreement that impacts collecting activity. Ethical regulations, such as the widespread adoption of Institutional Animal Care and Use Committees in the U.S. and similar bodies in other nations, add additional requirements to nascent biological collecting initiatives that must be properly administered by researchers. Rounding out regulatory hurdles facing biodiversity researchers and collectors are national and local regulations, which can range from non-existent to extremely restrictive. For example, in the U.S., the Lacey Act restricts the transport of certain taxa across state lines and adds import requirements in some contexts, restrictions that must be overcome for widespread collecting efforts in the U.S. These regulations are extremely important for a variety of ethical and conservation reasons and in many cases were adopted to combat harmful practices of commercial or black-market exploitation, but scientific research and collecting has also been impacted. While these international agreements clearly recognized the importance of scientific research on biodiversity, individual signatory nations may not recognize this intention, potentially stifling biodiversity research and collecting (Hamer et al. 2021). Finally, the landscape of research interests, funding, personnel, and scientific or taxonomic expertise is constantly shifting. Each of these factors, which play out across individual institutions and globally, may help to explain declines in collecting activity since the period of greatest collecting activity in the latter half of the 20th century. Individually and in combination, these factors help to explain declines in biological collecting and government and non-governmental organizations must continue to evaluate ways in which scientific collecting can be revitalized.

Diverse options are available to overcome the concerning trends evident in my temporal analysis of natural history collections growth, and I will make several higher-level recommendations for how we can ensure sustainable growth of natural history collections. Natural history collection holdings continue to be digitized and amassed in data portals like GBIF, but until now, there has been no central dashboard summarizing natural history collecting activity across taxonomic groups, nations, or institutions. Concerted effort is needed to improve digitization of individual collections and aggregate these data, but more specifically, natural history museums must invest in infrastructure and personnel that allows them to informatically explore existing data, as I have done here (see also (Johnson et al. 2023)), to better guide data digitization and aggregation and new specimen and artifact collection. Armed with such real-time information, governments, funding agencies, and individual institutions should consider implementing collecting quotas or recommendations for collecting that ensures sustainable growth, which should be informed both by our understanding of biodiversity across different taxonomic clades and our knowledge of the geographic or taxonomic specializations of institutions or nations. Today, most collecting activity is driven by the interests of individual curators and the funding they secure to pursue individual research programs. However, natural history collections are emerging as critical infrastructure for diverse areas of research beyond systematic investigations of the tree of life, which has historically been the focus of many museum practitioners. This trend, plus knowledge of best practices in experimental design that dictate the utility of natural history collections for experimental science, strongly argues for prescribed minimum collecting activity that is coordinated across various hierarchical levels. Fortunately, the recent announcement of required specimen management plans when applying for research grants from the U.S. National Science Foundation (see https://www.nsf.gov/pubs/2023/nsf23578/nsf23578.htm) may encourage more consistent collecting and deposition of specimens into natural history museums. Finally, funding levels for natural history collections need to be increased to ensure continued curation of existing holdings and sustainable growth through new collecting initiatives. While my data and analyses are insufficient to assess whether funding levels and priorities have contributed to diminished collections growth, documented closures and personnel layoffs (Dunnum et al. 2018) provide anecdotal evidence that financial austerity has negatively impacted natural history museums in recent years. Without a recommitment of funding agencies (governmental and philanthropic) and individual institutions to the mission of natural history museums and biorepositories through increased and sustainable funding, we risk diminished utility of natural history collections at a time where they are increasingly valuable for the scientific enterprise in the life sciences and beyond.

## Conclusions

This study presents the first detailed investigation of temporal patterns of global zoological natural history collections growth between 1900 and 2015. Collecting activity varied greatly over time, both globally and based on individual patterns of collections, nations, and taxonomic clades, and peaked around 2000, with recent years characterized by precipitous declines in collections growth. I also released an interactive data dashboard built using R Shiny to allow natural history museum stakeholders to explore these historical collections growth patterns in greater detail. In response to recent downward trends in collecting activity, I recommend that (1) institutions and other stakeholders invest more in informatic exploration of natural history museum holdings to guide future collecting, digitization, and aggregation efforts, (2) governments, funding agencies, and individual institutions implement collecting quotas or recommendations to ensure consistent, sustainable collecting activity in the future, and (3) government and philanthropic funding agencies increase financial commitments to natural history collections to catalyze more consistent future collections growth.

## Acknowledgements

I am grateful for constructive comments from Scott Edwards, Michaël Nicolaï, and two reviewers recruited by the journal: Gustav Paulay and Gary Rosenberg. I also acknowledge Vanya Rohwer and Matt Strimas-Mackey for advice on gathering natural history collection data from GBIF and Rhett Rautsaw for advice on building an interactive data dashboard in R Shiny. Postdoctoral support for D.C.C. has come from a fellowship from the United States National Science Foundation (DEB-1812310). The computations in this paper were run on the FASRC Cannon cluster supported by the FAS Division of Science Research Computing Group at Harvard University.

## Notes

https://github.com/darencard/NHMinformatics

https://doi.org/10.5281/zenodo.8393331

### Competing Interest Statement

The authors have declared no competing interest.

### Summary of Updates

Based on feedback from reviewers, I have modified the manuscript to better emphasize the full results, including the nuanced patterns of collecting activity, and place less emphasis on the finding of diminishing collecting growth, given that several biases could confound this result. I have included additional discussion sections noting the biases of the data and the analysis performed and discussing reasons for the apparent decline in collecting activity in recent years. I also have made small copy edits and followed several suggestions by the reviewers for improving readability.

